# Earl Grey ParTEA: scalable, consistent transposable element annotation from pangenomes to deep phylogenetic scopes

**DOI:** 10.64898/2026.11.04.764261

**Authors:** Tobias Baril, Alice Feurtey, Daniel Croll

**Affiliations:** Laboratory of Evolutionary Genetics, Institute of Biology, University of Neuchâtel, Rue Émile-Argand 11, 2000 Neuchâtel, Switzerland; Department of Bioinformatics and Genetics, Swedish Museum of Natural History, Stockholm, Sweden; Science for Life Laboratory (SciLifeLab), Swedish Museum of Natural History, Stockholm, Sweden

**Keywords:** transposon, repeat, mobile genetic element, genome, software, pipeline

## Abstract

Transposable elements (TEs) are major drivers of genomic diversity, yet understanding their mobility and functional impact requires their accurate description in genomes. TE annotation is still typically performed genome-by-genome. TEs are often shared between some members of the same or closely related species. Hence, TE annotation with a library from a single reference misses families absent from that assembly, whereas building independent per-genome libraries fails to consistently surface shared TEs, hindering comparison. Here, we present Earl Grey ParTEA, a fully automated Snakemake pipeline extending the functionality of Earl Grey for comparative TE annotation across genome panels. ParTEA builds a single non-redundant TE library from all input genomes, resolves chimeric consensus sequences, and uses this shared library to annotate every genome, enabling direct comparison of TE family diversity, abundance, shared and private content, and activity landscapes. Optional modules provide TE library saturation analysis, BUSCO-based assembly quality control, and phylogenetic ordering of outputs. We applied ParTEA to sets of genomes spanning evolutionary scales: ten *Arabidopsis thaliana* accessions, ten *Drosophila* species, and ten species spanning the fungal kingdom. The *Drosophila* TE library expanded from 343 to 1,852 curated families including a previously missed rolling circle element with over 6,000 copies in *D. miranda*. The proportion of TE families shared across all genomes was highest within *A. thaliana* (97.3%), 37.7% across *Drosophila* species and only 0.5% across the set of fungi. ParTEA is available through Bioconda and represents a computationally efficient reproducible framework for comparative TE analysis in the era of large-scale genome analyses.

## Introduction

Advances in sequencing technologies, coupled with declining costs, have led to a substantial increase in the availability of high-quality genome assemblies. These developments have shifted research beyond the analysis of single reference genomes towards the interrogation of intraspecific variation using large panels of genome assemblies (including pangenomes). Comparative analyses within species and genera have revealed extensive genomic variation, much of which is associated with variable transposable element (TE) content (Ǫuadrana et al. 2016; Baril and Hayward 2022; Rech et al. 2022; Baril et al. 2025; Ǫian et al. 2025).

TEs are mobile DNA sequences capable of moving within and between host genomes and can follow evolutionary trajectories partly distinct from those of their hosts (McClintock 1956). Most TE insertions are expected to range from nearly neutral to highly deleterious, reflecting their potential effects on genome integrity and gene regulation (Bourque et al. 2018). Consequently, hosts have evolved diverse mechanisms to suppress TE activity (Luo and Lu 2017; Schrader and Schmitz 2018; Lorrain et al. 2021), while deleterious insertions may also be removed by purifying selection (Bourque et al. 2018; Langmüller et al. 2023). Across populations, TE insertion polymorphisms are shaped by migration, drift, and selection (Bourgeois and Boissinot 2019). TE activity has therefore been proposed as an important source of genomic variation that, in some instances, could act over short evolutionary timescales and facilitate local adaptation (Tahami et al. 2024; Baril et al. 2025). Understanding these processes requires detailed intraspecific characterisation of TE insertions, while comparisons among related species may also illuminate mechanisms contributing to adaptation, the evolution of genome architecture, speciation, and the maintenance of biodiversity (Jurka et al. 2007; Jurka et al. 2011; Belyayev 2014).

The availability of high-quality genome assemblies (*i.e.* pangenomes) for an increasing number of non-model species provides substantial scope for comparative TE analyses. However, this also creates a substantial challenge because curated TE reference resources remain unavailable for much of the tree of life. As a result, studies commonly rely on *de novo* TE detection strategies that either (i) generate a TE library from a single genome assembly and use it to annotate all genomes, or (ii) generate separate TE libraries for each assembly and annotate genomes independently. Both strategies have important limitations. In the first case, TE families absent from the focal assembly cannot be detected. In the second, homologous TE families identified in different assemblies are difficult to reconcile across libraries, and shared elements are often redundantly represented under different names.

The availability of multiple genome assemblies per species, together with denser sampling across clades, is increasing rapidly. However, these advances have not been matched by the development of tools able to fully exploit such datasets. There is therefore a need for robust, scalable approaches that can analyse multiple assemblies while generating non-redundant and directly comparable TE annotations across genome assembly panels. Equally important is usability, as complex installation requirements, configuration burdens, and incompatible output formats can limit adoption and downstream reproducibility.

To address these challenges, we present Earl Grey **Pa**ngenome **T**ransposable **E**lement **A**nalysis (**ParTEA**), a fully automated pipeline for TE curation and comparative analysis that supports scalable investigation of TE diversity, turnover, and evolutionary dynamics across any number of genome assemblies at both intra- and inter-species scales. ParTEA combines assembly quality assessment with generation of non-redundant TE annotations, processing of chimeric consensus sequences, identification of shared and lineage-specific elements, and phylogenetically informed comparative summaries within and among species groups. Furthermore, we demonstrate how the ParTEA framework performs across three scales of comparative genomics: within species (*i.e.* pangenomes), using 10 *Arabidopsis thaliana* genome assemblies; within genera, using 10 *Drosophila* reference genome assemblies; and across a kingdom, using 10 fungal reference genome assemblies. The automation and highly efficient ParTEA pipeline integration standardises the analysis of multiple genome assemblies, enables consistent and comparable characterisation of TE landscapes across datasets spanning diverse levels of evolutionary divergence. Together, these capabilities provide a scalable framework for comparative TE analysis and facilitate the study of TE diversity across the tree of life.

## Implementation

ParTEA is a Snakemake-based pipeline that extends EarlGrey (Baril et al. 2024) to perform *de novo* transposable element (TE) library construction, annotation, and comparative analysis across multiple genomes simultaneously. To make ParTEA user-friendly, the package is provided in Bioconda (Grüning et al. 2018), and can also be installed directly from the GitHub repository (https://github.com/TobyBaril/EarlGreyParTEA). It can be run on Linux distributions, such as Ubuntu, and installed on local systems or HPCs supporting Conda package management. Implemented in Snakemake (Mölder et al. 2021), all steps are parallelised across input genomes where possible, with automated thread allocation so that analyses proceed concurrently. ParTEA can be run on HPCs dispatching the workload to SLURM or LSF queuing systems. The ParTEA pipeline is agnostic to the installed Earl Grey version, enabling users to install ParTEA alongside their current Earl Grey installations, but it is designed to work best with Earl Grey v7.3.0 onwards, to ensure compatibility with the latest implementations of the Dfam database (v4.0)(Hubley et al. 2016; Storer et al. 2021).

ParTEA runs in a configured environment to avoid conflicts between tool versions and to streamline the installation procedure. Full installation instructions are provided in the GitHub repository and minimal configuration is required by the user, with the exception of specifying which partitions of the Dfam database (Hubley et al. 2016; Storer et al. 2021) the user would like to configure the FamDB (v3.0.0) tool with. The instructions to do this are automatically generated by ParTEA, which detects whether Dfam partitions have been configured previously.

Once installed, ParTEA will run on all the genome assemblies provided in the configuration file, which serves as the central location where all user-specified options are stored. After conda environment activation, ParTEA can be called with one of three commands: ‘earlGreyParTEÀ (full pipeline run), ‘earlGreyParTEA _LibConstruct’ (only run up to the creation of the TE library), or ‘earlGreyParTEA_AnnotationOnly’ (run annotation and post-processing using a user-supplied TE library). Editable example configuration files can be generated from any of the three commands, with automated configuration file generation possible with appropriate flag usage (Table 1).

**Table 1.** Parameters for ParTEA. FILE=path to file; INT=integer value; DIR=path to directory; PART=slurm partition/LSF queue; ACCT=slurm account/LSF project; FLAGS=other flags for specific tools.

| Command Flag | Description | Required? |
|---|---|---|
| -c, --config [FILE] | Path to config.yaml file containing all options to run ParTEA | Y |
| -t, --threads [INT] | Number of threads to use<br>Local run: total threads available<br>SLURM/LSF run: max threads per job | Y |
| -m, --memory [INT] | Scheduling memory budget (Snakemake will not start new jobs if doing so would exceed this budget) | N |
| -n, --dry-run | Perform dry run to check what would be executed | N |
| -h, --help | Show help message | N |
| **Config File Generation** |  |  |
| --generate-config [FILE] | Generate a template config.yaml file | N |
| --genome-dir [DIR] | Used with --generate-config, the config.yaml will be populated with all genome assemblies found in the specified directory | N |
| --from-csv [FILE] | Used with --generate-config, the config.yaml will be populated with genome assemblies and names from a 2-column csv (species, genome_path) | N |
| --output-dir [DIR] | Used with --generate-config to specify the desired output directory | N |
| **SLURM Options** |  |  |
| --slurm | Submit jobs to SLURM cluster | N |
| --slurm-partition [PART] | SLURM partition/queue that jobs will be submitted to. This can also be set in the config file. | N |
| --slurm-account [ACCT] | SLURM account string. This can also be set in the config file. | N |
| --slurm-jobs [INT] | Max number of concurrent jobs to submit to SLURM.<br>Default = genomes x 3, capped at 200. | N |
| --slurm-extra [FLAGS] | Additional flags that users specify with sbatch. | N |
| **LSF Options** |  |  |
| --lsf | Submit jobs to LSF cluster | N |
| --lsf-jobs [INT] | Max number of concurrent jobs to submit to LSF.<br>Default = genomes x 3, capped at 200. | N |
| --lsf-queue [PART] | LSF partition/queue that jobs will be submitted to. This can also be set in the config file. | N |
| --lsf-project [ACCT] | LSF project string. This can also be set in the config file. | N |
| --lsf-extra [FLAGS] | Additional flags that users specify with bsub. | N |
| **Snakemake Options** |  |  |
| --unlock | Unlock working directory after a crash | N |
| --rerun-incomplete | Rerun incomplete jobs from a previous run | N |
| --snakemake-args [FLAGS] | Additional Snakemake arguments | N |

Users can set pipeline behaviour from a central point, using the generated yaml file. Here, the user can control all optional aspects of the pipeline, with behaviour consistent with Earl Grey. Config files used in analysis are provided in supplementary file 1. In addition, several new ParTEA-specific features have been developed: (i) Analysis of shared vs unique TEs; (ii) user-specified redundancy reduction in TE consensus libraries, consistent renaming, and resolution of chimeric consensus sequences; (iii) Analysis of TE diversity saturation (*i.e.* how many genomes were required to detect all TEs present in the analysed genome assemblies); (iv) BUSCO annotation; (v) Basic phylogenetic reconstruction using supermatrices of BUSCO single-copy orthologs; (vi) Presentation of all analyses in phylogenetic order (based on BUSCO supermatrix phylogenetic tree). The steps involved in the ParTEA procedure, including optional analyses, are outlined below. Unless stated, workflows are implemented using the Earl Grey dependency, and thus function as previously described (Baril et al. 2024):

1. **Workflow visualisation:** Directed acyclic graph (DAG) visualisations of the complete Snakemake workflow are generated at run start using Graphviz in SVG format, providing a visual record of the exact set of rules and dependencies executed for a given configuration. An overview of all processes is provided in figure 1.
2. **Genome preparation (rules: prep_genome):** Each input genome is first prepared by stripping whitespace from FASTA headers, replacing ambiguous IUPAC nucleotide codes (D, V, H, B, P, E) with N, and renaming sequences using a short internal key via the EarlGrey ‘headSwap.sh’ utility. A gzip-compressed backup of the original genome and a sequence dictionary are retained alongside the prepared file for downstream use.
3. **Optional - initial repeat masking (rules: repeatmasker_warmup, repeatmasker, repeatmasker_custom, extract_repeatmasker_library):** If a RepeatMasker species clade (repeatmasker_species) or a custom TE library (custom_library) is provided in the configuration, each prepared genome is masked in a preliminary RepeatMasker (v4.2.4) pass prior to *de novo* discovery. This reduces the proportion of known repeats that RepeatModeler2 (Flynn et al. 2020) must model independently and retains known elements. A single warmup job pre-builds the RepeatMasker BLAST cache before parallel genome jobs begin, preventing race conditions on first use. In addition, if a RepeatMasker species clade is specified, a copy of the TE library is extracted from the local Dfam database and saved in FASTA format. This library is then used in two independent ways: (a) for the initial masking pass, and (b) it is added to the combined pre-clustering library with a REPMASKER_ prefix.
4. ***De novo* TE discovery (rules: build_db, repeatmodeler):** A BLAST database is constructed from the (optionally masked) genome using the BuildDatabase function of RepeatModeler2 (v2.0.9). RepeatModeler2 is then run on this database to perform *de novo* identification of repetitive sequence families, generating a consensus library of candidate TE sequences for each input genome assembly.
5. **Library extension and refinement (rules: testrainer):** The RepeatModeler consensus library for each genome is passed to the TEstrainer module of Earl Grey, which iteratively extends and refines each consensus sequence through a configurable number of BLAST-extend-align-trim cycles (default: 10 iterations). In each cycle, each consensus is used to search the genome by BLAST (v2.17.1+), flanking sequence is extracted, aligned, and trimmed, and an updated consensus is derived. Sequences supported by fewer than the minimum number of aligned copies (min_consensus_seqs; default: 3) are discarded, and the final refined library is output as a strained FASTA file.
6. **ParTEA feature - combined TE library construction (rules: cluster_all_species, split_chimeras):** The per-genome strained libraries are concatenated, with each sequence header prefixed by the species identifier to ensure traceability. If a RepeatMasker clade library or custom library was provided, these are also included (prefixed REPMASKER_ and CUSTOM_, respectively). By default, the combined library is clustered with cd-hit-est (v4.8.1)(Li and Godzik 2006; Fu et al. 2012) to produce a single non-redundant TE library (combined_all_species.clstrd.fa). Four clustering parameters are configurable: sequence identity (clustering_identity, default: 0.8, equivalent to ‘-c’ in cd-hit-est), short-sequence alignment coverage (clustering_coverage, default: 0.8, equivalent to ‘-aS’ in cd-hit-est), long-sequence alignment coverage (clustering_coverage_long, default: 0.0 (no limit), equivalent to ‘-aL’ in cd-hit-est), and minimum length ratio between the shorter and longer sequence (clustering_length_diff, default: 0.5, equivalent to ‘-s’ in cd-hit-est). The identity and coverage defaults implement the pragmatic 80-80-80 rule (Wicker et al. 2007; Goubert et al. 2022). We note that the 80-80-80 rule provides a convenient but imperfect approximation of TE family structure, as it may over-merge young families or over-split ancient or highly diverged families. In addition, the rule can perform poorly for certain elements such as DIRS-like elements and non-LTR retrotransposons. Setting clustering_coverage_long to a value such as 0.75 is recommended when the combined library contains long sequences that could otherwise act as spurious cluster hubs, absorbing shorter unrelated members. Clustering can be disabled (skip_clustering: true), in which case all sequences are retained and concatenated directly into the final library. Optionally, a post-clustering chimera detection step can be enabled (split_chimeras: true). Chimeric cluster representatives arise when consensus building joins sequences from two or more distinct TE families into a single sequence (i.e. the chimera). To resolve these cases, the detection algorithm proceeds as follows. For each cluster in the cd-hit-est .clstr file, the pairwise alignment coordinates that non-representative members reported against the cluster representative are extracted. An overlap graph is then constructed in which two members are connected by an edge if their alignment windows on the representative overlap by at least chimera_overlap_min nucleotides (default: 50). Connected components of this graph are identified by breadth-first search. A cluster is flagged as chimeric if it yields ≥2 components, each spanning at least chimera_min_component_span of the representative’s length (default: 10%), and the cluster contains at least chimera_min_members non-representative members (default: 3; note that clusters with fewer members may lack the statistical power to distinguish genuine chimerism from partial or low-coverage alignments). For each confirmed chimeric cluster, the original representative is retained in the output with a _CHIMERA suffix for traceability, and each component is represented by its longest member sequence, retrieved from the pre-clustering combined FASTA to preserve the original species-prefixed header and TE classification tag. The resulting ‘combined_all_species.chimera_split.fà replaces the standard clustered library as the downstream input when chimera splitting is enabled. A tab-separated summary (chimera_detection_summary.tsv) is written recording, for each cluster, whether it was flagged as chimeric, the number of components detected, component sizes, and a chimera score. To quantify the degree of chimerism, we define a chimera score for each flagged cluster as the largest gap between adjacent alignment component regions, normalised by the length of the cluster representative. Formally, for a representative of length *L* with *k* validated components, the chimera score is:

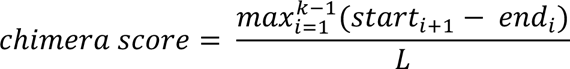 Where start_i+1_ and end_i_ are the leftmost and rightmost alignment coordinates of adjacent components on the representative, respectively. The score ranges from 0 to 1, where values approaching 1 indicate that the two putative TE families are cleanly separated by a large unaligned region, consistent with a head-to-tail fusion of two distinct consensuses, whilst values near 0 indicate that the two component regions meet at nearly the same position on the representative, reflecting a more ambiguous boundary.
7. **ParTEA feature - TE family saturation analysis (rules: saturation_plot):** A saturation curve is generated from the cd-hit-est cluster membership file to assess whether sampling of TE diversity is approaching saturation across the input genome set. The analysis permutes the order in which genomes are added to the analysis (default: 100 permutations) and records the cumulative number of unique TE families (clusters) after each addition. The mean and 95% confidence interval across permutations are plotted as a curve (saturation_plot.pdf) and accompanying tabular data are written to saturation_data.tsv. If clustering of library sequences is skipped, these plots are generated based on the number of families discovered in each genome, but no deduplication is performed, which may lead to elevated TE family counts. If a RepeatMasker clade library or custom library was provided, the number of sequences from this set serves as the baseline above which new families per genome are shown.
8. **Dataset-wide TE annotation (rules: repeatmasker_annotation):** Each prepared genome is annotated using RepeatMasker with the combined ParTEA TE library as the query. RepeatMasker is invoked with the options as specified in Earl Grey, namely a sensitive search with the exclusion of insertion sequences (-s -no_is). The resulting .out and .tbl files are used in all subsequent annotation processing steps.
9. **Helitron detection (optional) (rules: heliano_detection):** If Helitron detection is enabled (run_heliano: true), HELIANO (v1.3.1)(Li et al. 2024) is run on each genome in parallel with RepeatMasker. HELIANO detects rolling circle elements (also termed *Helitrons*). Predicted Helitron loci are converted to GFF3 format and merged with RepeatMasker outputs in the subsequent step.
10. **Repeat merging, filtering, and summarisation (rules: merge_repeats, generate_summary_charts):** RepeatMasker output coordinates and, where available, HELIANO predictions are merged and filtered using the EarlGrey rcMergeRepeats utility. Consistent with optional settings in Earl Grey, if margin: true is set in the config file, short TEs (<100bp) are removed from the final annotations. If loose merging fails for a genome, a strict-merge fallback is attempted automatically. The final annotations for each genome are written to BED and GFF3 files. A high-level count of TE classes and a summary pie chart are generated using the EarlGrey autoPie.sh utility.
11. **Divergence estimation and repeat landscapes (rules: calculate_divergence):** Kimura two-parameter (K2P) divergence between each annotated TE copy and its consensus is calculated using the EarlGrey divergence_calc.py script. Divergence values are appended to the GFF3 annotation file and a repeat landscape (divergence histogram) is plotted per genome using the EarlGrey divergence_plot.R script. A tabular divergence summary is written to the per-genome summary directory.
12. **Output consolidation (rules: sweep_up_files, generate_softmasked_genome):** Final annotation BED and GFF3 files are copied into a per-genome directory alongside all summary charts, divergence plots, and landscape plots, providing a self-contained result set for each genome, with the same contents as if an individual Earl Grey run was performed on each genome. If softmasking is requested (softmask: true), a softmasked version of the original genome is produced using BEDTools (v2.31.1)(Ǫuinlan and Hall 2010) maskfasta.
13. **ParTEA feature - Shared and unique TE content analysis (rules: shared_unique_plot, shared_unique_plot_phylo):** When run_shared_unique: true is set, the pipeline computes the proportion of TE families and genome coverage that is shared across all genomes, partially shared among subsets, or private to individual genomes. In full pipeline mode, cluster membership from the cd-hit-est run is used to assign a consistent TE family identity to sequences across genomes, so that homologous families are treated as the same family even where per-genome naming differs. Whether a family is classified as shared or private is then determined by its annotation in each genome (*i.e.* whether it has ≥1 hit in that genome’s GFF). Results are presented as stacked bar charts for both TE family counts and genome coverage fractions (shared_unique_families.pdf, shared_unique_coverage.pdf), with companion TSV files for downstream use. In addition, a heatmap is automatically produced to show TE coverage across all input genomes for the most abundant TE families. When the BUSCO phylogenomics module is also active, phylogenetically ordered versions of these charts are also produced.
14. **ParTEA feature - BUSCO quality control & phylogenomic tree (rules: fetch_busco_db, run_busco, busco_summary_table, extract_busco_aa, align_b usco_gene, create_supermatrix, run_fasttree, busco_completeness_phylo, bus co_te_qc):** When run_busco_phylo: true is set, a lightweight phylogenomic inference pipeline is executed alongside the main annotation workflow. BUSCO (v6.1.0)(Simão et al. 2015; Manni, Berkeley, Seppey, and Zdobnov 2021; Manni, Berkeley, Seppey, Simão, et al. 2021) is run in genome mode on each prepared genome using a user-specified lineage dataset (busco_lineage). A TSV is generated recording the occupany of each candidate gene, and single-copy orthologue amino acid sequences that are present in at least a configurable fraction of genomes (busco_min_occupancy; default: 0.5) are extracted and independently aligned with MAFFT (v7.526)(Katoh and Standley 2013), with poorly aligned columns removed by ClipKIT smart-gap (v2.12.0)(Steenwyk et al. 2020). Trimmed alignments are concatenated into a supermatrix using PhyKIT (v2.1.90)(Steenwyk et al. 2021), and a maximum-likelihood species tree is inferred using FastTree (v2.2.0)(Price et al. 2009) with the Le-Gascuel 2008 model (-lg) and length rescaling to optimise the Gamma20 likelihood (-gamma). When both run_busco_phylo and run_shared_unique are enabled, a stacked BUSCO completeness bar chart ordered along the inferred cladogram is generated, as is a scatter plot of BUSCO completeness against TE genome coverage per species to aid quality control of the input assemblies.

**Figure 1.**
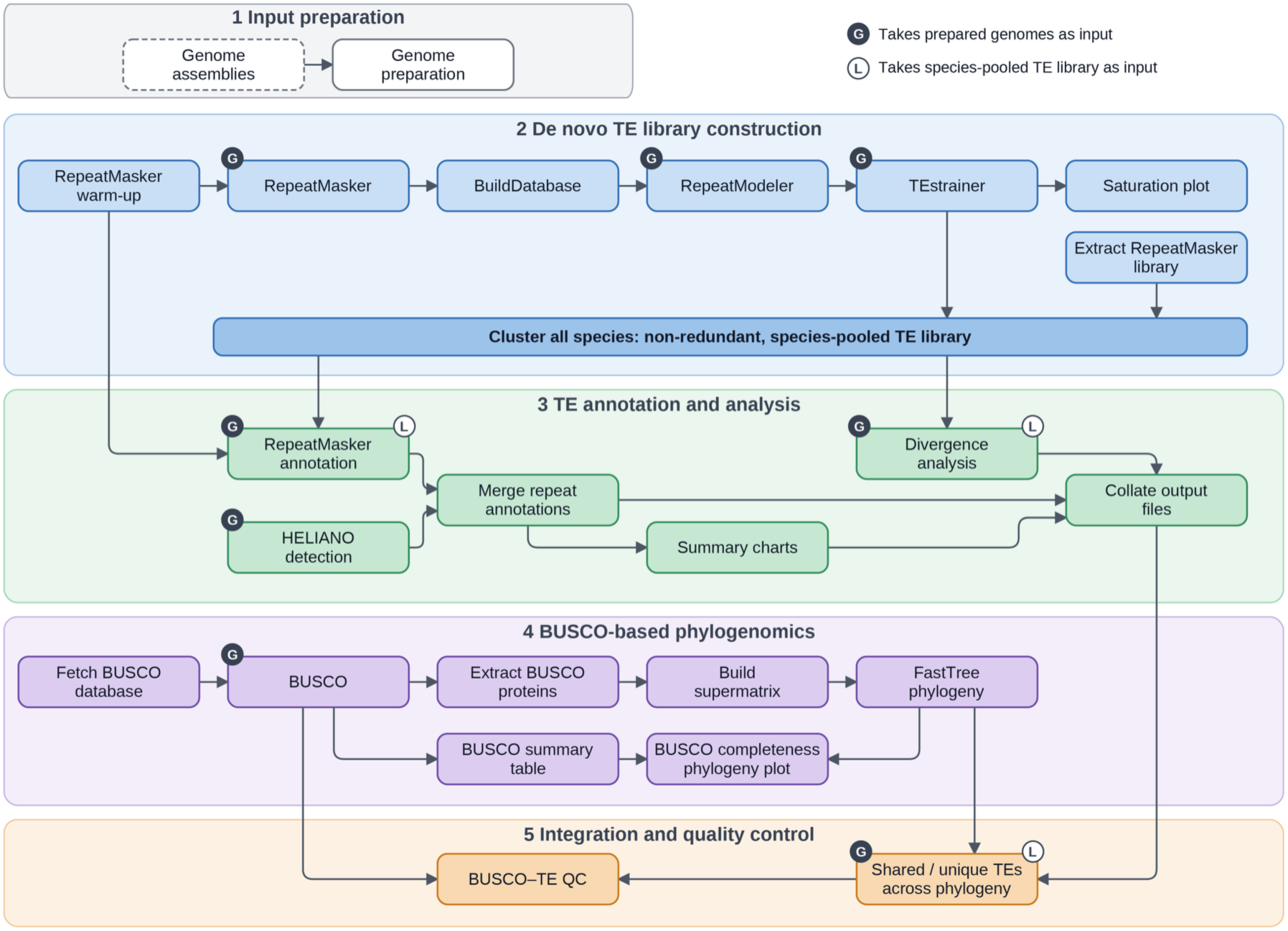
Simplified overview of the Earl Grey ParTEA annotation pipeline. Steps are partitioned into functional groups performing specific stages of analysis, as labelled. Boxes 4 and 5 indicate processes that run when the optional BUSCO analysis is implemented. **ALT TEXT:** Flow chart summarising the processing steps for Earl Grey ParTEA. Subsections are labelled from 1 to 5, illustrating steps forming core stages of the pipeline.

As the underlying workflow utilises the existing Earl Grey pipeline, analyses are backwards-compatible with those perviously performed.

## Materials and Methods

To demonstrate the new features of ParTEA, and to assess the impact of consistent panel-wide TE analyses, we analysed three publicly available datasets. We obtained 10 long-read sequencing assemblies of *Arabidopsis thaliana* from a recent pangenome analysis (Igolkina et al. 2025), 10 reference genome assemblies for the *Drosophila* genus from NCBI, and 10 reference genome assemblies spanning the fungal kingdom (Table S1, supplementary file 3). In all three phylogenetic scopes, all optional ParTEA processes were activated and intial RepeatMasker runs were specified with species terms “arabidopsis”, “7215” (the NCBI taxonomy ID for the *Drosophila* genus), and “fungi” respectively. For the BUSCO assessments, brassicales_odb12 was used for *A. thaliana*, drosophila_odb12 was used for *Drosophila*, and fungi_odb12 was used for fungi. The minimum occupancy for a BUSCO gene to be included in the supermatrix for phylogenetic reconstruction was set to 5% of the total number of assemblies (busco_min_occupany: 0.05). The pipeline was invoked using the slurm options to make use of the HPC queueing system and fairshare policy. Config files for each analysis are provided in supplementary file 1. Complete output directories for each analysis are provided in supplementary file 2.

### Synteny Analysis

We used NGenomeSyn (v1.50)(He et al. 2023) to examine the genomic distribution of TE family *GCF_003369915_1_rnd-1_family-397* across Drosophila genomes. Pairwise genome assemblies were aligned in phylogenetic order, as determined by the ParTEA phylogeny subroutine, using MiniMap2 “-x asm5” (v2.31-r1302)(Li 2018). PAF alignments were converted to NGenomeSyn link files using the Paf2Link utility, retaining alignments with at least 5,000 matching bases. Alignments for adjacent genome pairs were used to draw ribbons between neighbouring tracks. Short contigs <1Mb were removed to improve visualisation. NGenomeSyn was used to plot the genomes in phylogenetic order, with bed files containing locations of *GCF_003369915_1_rnd-1_family-397* provided as “special genome features” to enable plotting of their genomic distributions. Following plot generation, short contigs in *D. ananassae* with no links among genomes were removed to improve readability.

## Results

### Within-Species Application: Arabidopsis

To demonstrate the application of ParTEA within a single species or a pangenome context, we used ten chromosome-level genome assemblies for *A. thaliana* (Igolkina et al. 2025) (Supplementary Table S1). The ParTEA run completed in a total of 15 hours on a HPC system, with each job able to use up to a maximum of 16 threads. All *A. thaliana* assemblies show BUSCO single-copy completeness levels ≥98.7% (Figure 2).

**Figure 2.**
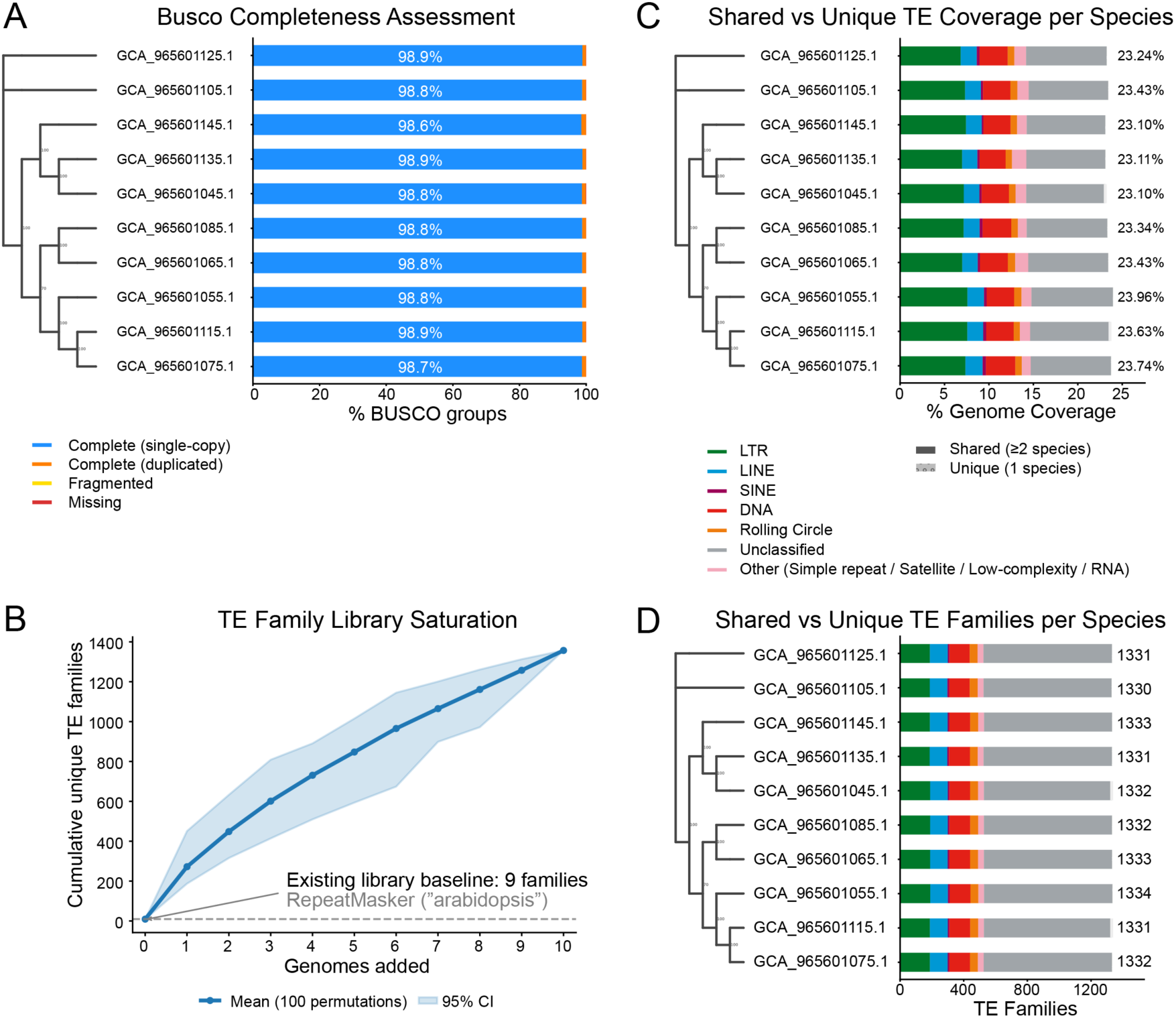
ParTEA output visualisations for Arabidopsis thaliana. All plots are generated automatically by ParTEA. **(A)** BUSCO completeness with samples in phylogenetic order as determined by the ParTEA supermatrix-based phylogeny assessment. **(B)** Saturation curve illustrating the impact of adding additional genomes to overall TE library composition. Confidence intervals are determined using a permutation approach with the number defined by the user (default: 100 permutations). **(C)** Shared and unique TE coverage (% of total genome size) for each assembly in the focal genome panel. Samples are arranged in phylogenetic order. Solid bars represent TE content arising from families shared by at least two genome assemblies, whilst faded spot bars represent TE content arising from TE families unique to the focal genome assembly. TE classifications are indicated by the key. **(D)** Shared and unique TE family count for each assembly in the focal genome panel. Samples are arranged in phylogenetic order. Solid bars represent TE family counts for families shared by at least two genome assemblies, whilst faded spot bars represent TE family counts for TE families unique to the focal genome assembly. Numbers at the ends of bars indicate total TE family count within each genome assembly. **ALT TEXT:** Graphs and data on BUSCO completeness and transposable element content annotated among 10 Arabidopsis accessions, with subfigures labelled from a to d.

The addition of each extra genome assembly increases the number of unique TE families considerably, up to a total of 1,357 *de novo* detected TE families across all ten genome assemblies (Figure 2, Supplementary Table S2, Supplementary File 3). Saturation is clearly not reached, indicating that the ten genome assemblies analysed are not sufficient to detect all TE diversity within *A. thaliana* (Figure 2), consistent with levels of TE diversity described previously in a larger *A. thaliana* genome panel (Igolkina et al. 2025).

Annotated TE abundance is remarkably stable across all *A.* thaliana accessions, with a mean total genome coverage of 28.02Mb and little variation among individuals (minimum genome coverage = 27.5Mb, GCA_965601045.1; maximum genome coverage = 28.9Mb, GCA_965601055.1). Nearly all annotated TE families are shared among all ten accessions (97.3%, 1,309/1,345 families). The most dominant TE families are LTR retrotransposons, with the top five annotated TE families contributing a mean of 3.62Mb and 1,452 copies across the 10 accessions (Table S3, Supplementary File 3). Only two TE families were private and annotated in a single assembly: an LTR retrotransposon (*GCA_965601145_1_rnd-1_family-45*) was annotated only in accession GCA_965601145.1, and an unclassified TE (*GCA_965601075_1_rnd-1_family-154*) was annotated only in accession GCA_965601115.1.

The lack of private families is expected at the within-species scale, given the shared evolutionary history and short divergence time among individuals. Consistent with this, TE activity landscapes were broadly similar among *A. thaliana* accessions (Supplementary Figure 1), with all profiles showing a similar distribution and a pronounced peak at low Kimura 2-Parameter divergence from consensus sequences. This points to recent activity of TEs in all accessions with the main contributors being LTR retrotransposons and DNA transposons, with LINE activity occurring in the more distant past (Supplementary Figure 1).

### Within-Genus Application: Drosophila

To evaluate the ability of ParTEA to analyse genomes at a broader phylogenetic scale, we applied the pipeline to ten chromosome-level assemblies representing distinct species of the genus *Drosophila*. The analysis was completed in a total of 23 hours on a HPC system, with individual jobs able to use a maximum of 16 threads.

Assembly completeness was generally high across the *Drosophila* dataset, with BUSCO single-copy complete scores exceeding 97% for all assemblies except *Drosophila miranda*, which had 84.6% single-copy complete BUSCOs, although including duplicated BUSCOs (14.6%) leads to a total BUSCO completeness of 95.2%, suggesting that its lower single-copy completeness is associated primarily with BUSCO duplication rather than widespread loss or fragmentation (Figure 3).

**Figure 3.**
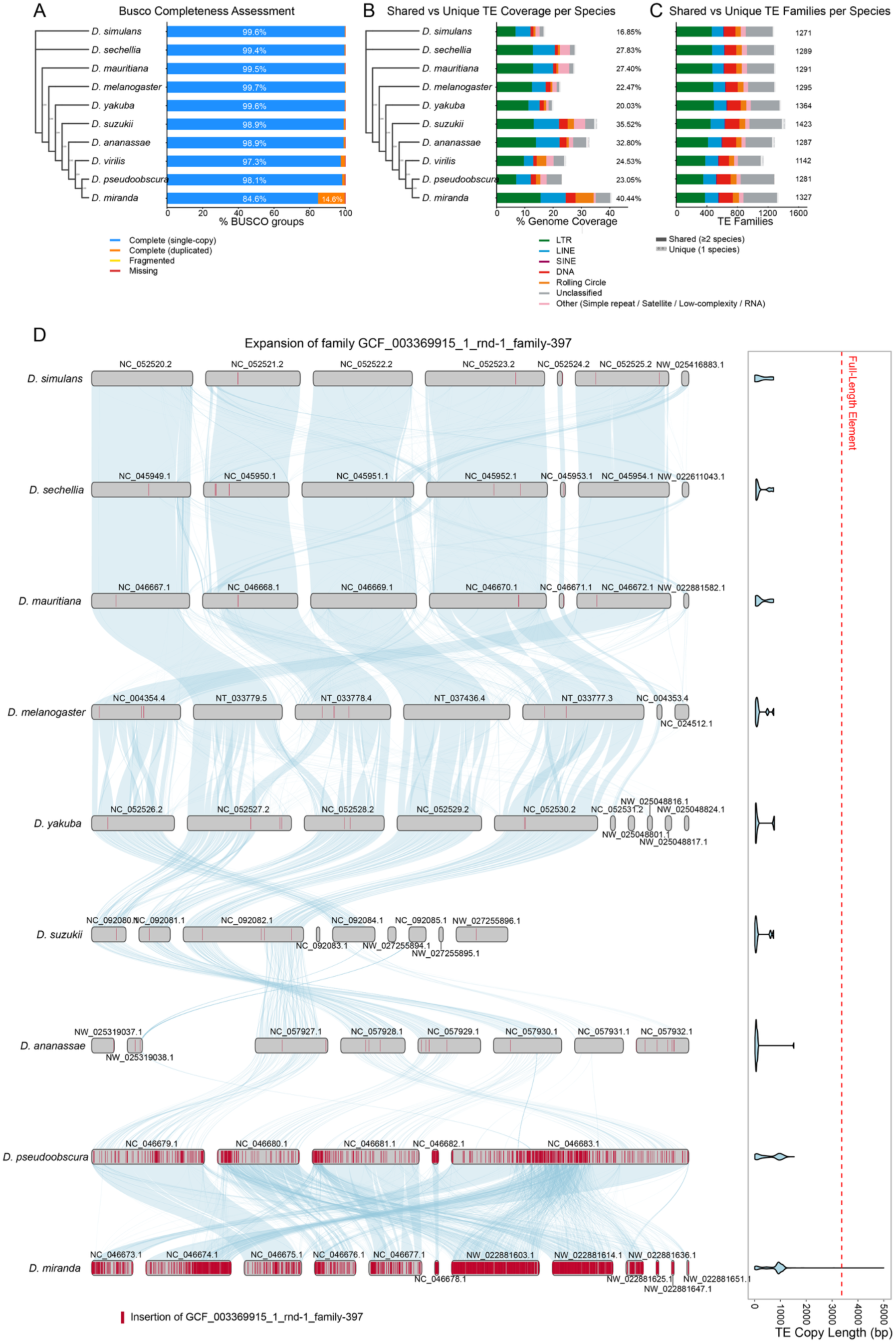
Earl Grey ParTEA annotation summaries across the Drosophila genome panel and distribution of TE family GCF_003369915_1_rnd-1_family-337. (A) BUSCO completeness, with assemblies ordered according to the ParTEA supermatrix-based phylogeny. (B) Shared and unique TE coverage, expressed as a percentage of assembly size. Solid bars show coverage from TE families detected in at least two assemblies; faded bars show coverage from families detected only in that assembly. Colours indicate TE classification, and end labels show total TE coverage. (C) Counts of shared and unique TE families, using the same definitions and colour key as in (B); end labels show the total number of TE families in each assembly. (D) Synteny among nine Drosophila species, ordered according to the ParTEA phylogeny. Grey bars represent chromosomes or scaffolds, ribbons show syntenic links between adjacent assemblies, and red marks indicate loci of GCF_003369915_1_rnd-1_family-397. Violin plots show the length distribution of annotated insertions; the dotted line marks the length of a full-length element. D. virilis was omitted from panel D to improve readability: after filtering, only 17 and 11 synteny links were retained in its comparisons with the adjacent species, consistent with its more divergent placement in the ParTEA phylogeny. **ALT TEXT:** Graphs on BUSCO completeness and transposable element content annotated among annotated Drosophila species, with subfigures labelled from a to d. Subfigures a to c show summaries of transposable element content. Subfigure d shows synteny among 9 Drosophila species as a graphical representation of the expansion of a single rolling-circle family.

Despite the large community TE library for *Drosophila*, comprising 343 families, the ParTEA cumulative TE library increased substantially as additional genomes were incorporated. The library increased to 1,852 families after inclusion of all ten genomes without evident saturation (Supplementary File 2). The accumulation of TE families demonstrates that substantial additional TE diversity can be identified when ParTEA is applied across multiple species, even within a single genus, and the absence of an obvious plateau further suggests that additional *Drosophila* genomes may lead to the discovery of currently unrepresented TE families.

The majority of TE families annotated among *Drosophila* (94.5%; 1,746 / 1,847 families with at least one annotation) are shared among at least two species (Table S4, Supplementary File 3), whilst a lower proportion of TE families are shared among all 10 *Drosophila* species (37.7%; 697 / 1,847 families with at least one annotation) than among all 10 *A. thaliana* accessions (97.3%, 1,309 / 1,345 families with at least one annotation), demonstrating the divergence in TE landscapes among species with increasing phylogenetic distance.

The most abundant TEs among *Drosophila* are a single LINE family (*DMCR1A*) and four LTR retrotransposon families (*Gypsy-8_DMel*, *Osvaldo8*, *Diver2*, and *GATE-1*) (Table S4, Supplementary File 3). These TE families are curated in Dfam 4.0, reflecting historic focus and effort to curate TEs in at least a subset of *Drosophila* species (*i.e. D. melanogaster*). Of note, however, is the rolling circle element *GCF_003369915_1_rnd-1_family-397*, which is found at low copy number (<50 copies) and low coverage (<5kbp) in eight of the ten species but has expanded to 1,024 copies (586kbp) in *D. pseudoobscura* and 6,145 copies (4.6Mb) in *D. miranda* (Figure 3D, Table S4, Supplementary File 3). This rolling circle element was only discovered during the *de* novo detection phase in the assembly of *D. miranda*, meaning this highly abundant component of *D. miranda* and *D. pseudoobscura* would have remained undetected if the existing Dfam *Drosophila* TE library had been used to annotate all genomes, or if *de novo* TE family detection had not included *D. miranda* in the analysis. We observe a reduction in synteny between *D. ananassae* and *D. pseudoobscura* co-occurring with the large increase in copy number and distribution of *GCF_003369915_1_rnd-1_family-397*, TK. Further, analysis of copy number lengths of *GCF_003369915_1_rnd-1_family-397* among the *Drosophila* genomes reveals no full-length insertions remain in any of the sampled genomes, with the exception of *D. miranda*. This suggests that ongoing activity of this TE family is limited given the lack of elements likely capable of transposition due to truncation, and that these expansions observed in *D. pseudoobscura* and *D. miranda* likely occurred in a burst of activity in the more distant past. This is further supported by the divergence of genomic copies of *GCF_003369915_1_rnd-1_family-397* in *D. pseudoobscura* (μ = 12.34%, SD = 8.87%) and *D. miranda* (μ = 7.43%, SD = 6.15%) (Supplementary File 2), indicating degradation of genomic copies following insertion. This example highlights the importance of incorporating a *de novo* approach in the annotation of genomes, especially in comparative genomics, where curated elements in existing libraries are unlikely to exhaustively cover extant TE diversity across the whole species set, leading to incomplete TE annotations and the hiding of genomic components that may be important contributors to genome architecture or differentiation among species.

At this broader genus-wide scale, the majority of TE content annotated among *Drosophila* is shared among at least 2 species, with a mean of 53.4Mb (99.2% of total TE content) coming from shared families. In contrast, a mean of 509kb (0.77% of total TE content) was annotated in a single genome, thus private TE families are rare among this sample of the *Drosophila* genus.

Together, these results demonstrate that application of ParTEA across multiple species within a single genus substantially expands the detectable TE repertoire while simultaneously resolving shared and lineage-specific components of TE diversity. The *Drosophila* analysis therefore illustrates how progressively incorporating additional genomes can improve characterisation of TE diversity beyond that captured by a single-species reference library and demonstrates the importance of such an integrative approach to more exhaustively characterise TE diversity within a genus.

### Within-Kingdom Application: Fungi

To demonstrate ParTEA’s ability to analyse genomes across large phylogenetic distances, we used ten genome assemblies spanning the fungal tree of life, including one assembly from each of the phyla: Cryptomycota, Microsporidia, Blastocladiomycota, Zoopagomycota, Mucromycota, and Chytridiomycota, and two assemblies from the phyla Basidiomycota and Ascomycota (Table S1, Supplementary File 3). The genome assemblies available with the highest scaffold N50 were selected for each phylum. The ParTEA run completed in a total of 39 hours on a HPC system, with each job able to use up to a maximum of 16 threads.

Genome assembly quality was more variable among the fungal genomes than either the *Arabidopsis* or *Drosophila* datasets (Table S1, Supplementary File 3), which could impact BUSCO completeness levels. In addition, Microsporidia and Cryptomycota have undergone extreme genome minimisation, leading to reduced BUSCO completeness when these lineages are analysed with the fungi database (Manni, Berkeley, Seppey, Simão, et al. 2021). Consequently, BUSCO completeness was highly variable across the fungi dataset, from 10.9% single-copy completeness in *Nematocida parisii* (Microsporidia) to 98.1% single-copy completeness in *Tricholoma hemisulphureum* (Basidiomycota) (Figure 4). The phylogenetic ordering implemented with sparse sampling across deep phylogenetic time has generated spurious placements, including *N. parisii* being placed between Ascomycota and Basidiomycota. This is likely driven by the large variance in BUSCO set completeness used to generate the phylogeny, and users should be aware of the potential for such artefacts in very homogeneous datasets spanning large evolutionary distances (Figure 4).

**Figure 4.**
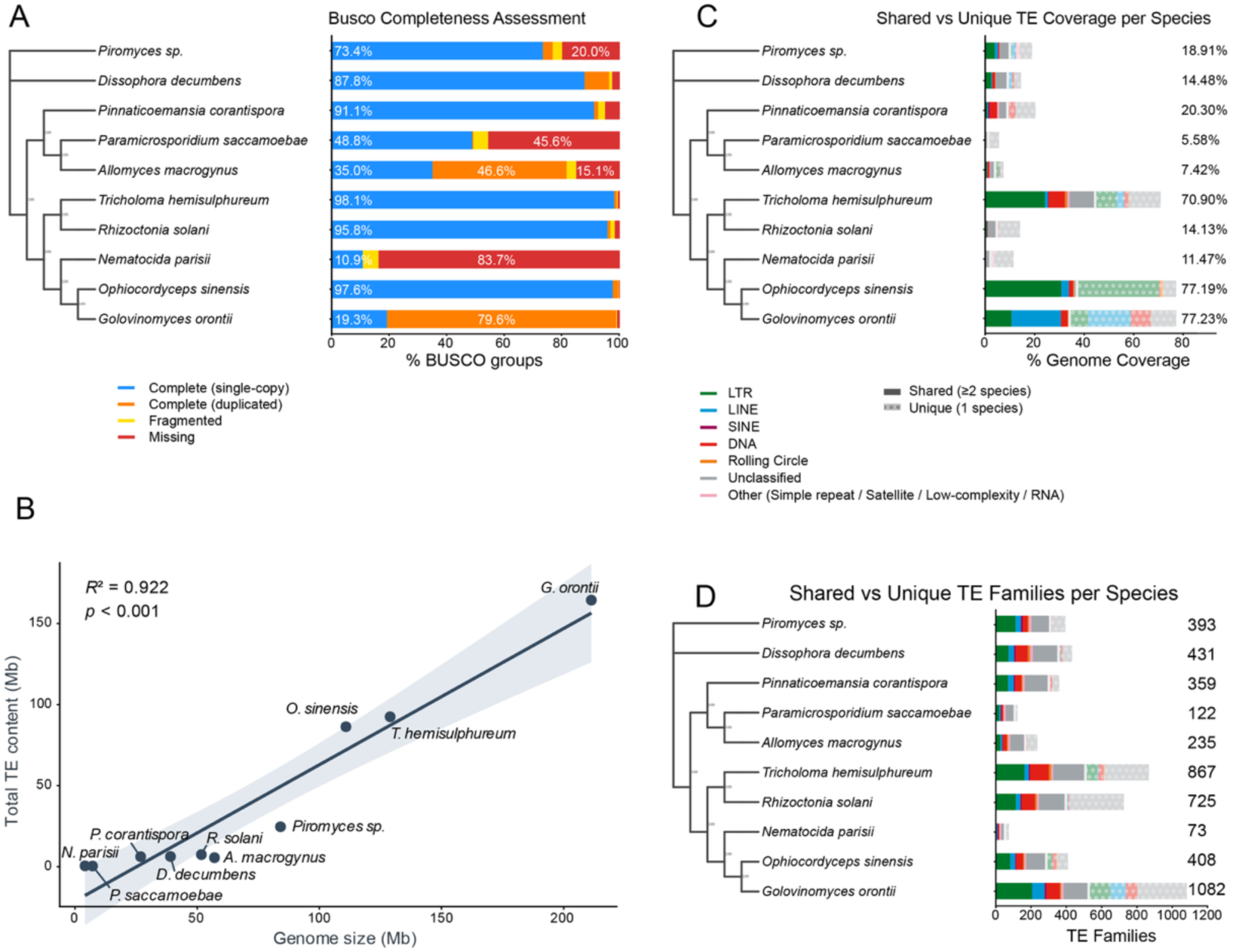
Earl Grey ParTEA-identified TE diversity among fungi. **(A)** BUSCO completeness with samples in phylogenetic order as determined by the ParTEA supermatrix-based phylogeny assessment. **(B)** Relationship between TE abundance and genome size across fungi, modelled using linear regression (Total TE content ∼ Genome Size). Shaded area indicates the S5% confidence interval. Model R^2^ and p value are provided in the plot. **(C)** Shared and unique TE coverage (% of total genome size) for each assembly in the focal genome panel. Samples are arranged in phylogenetic order. Solid bars represent TE content arising from families shared by at least two genome assemblies, whilst faded spot bars represent TE content arising from TE families unique to the focal genome assembly. TE classifications are indicated by the key. Numbers at the ends of bars indicate total TE coverage as a percentage of genome size within each genome assembly. **(D)** Shared and unique TE family count for each assembly in the focal genome panel. Samples are arranged in phylogenetic order. Solid bars represent TE family counts for families shared by at least two genome assemblies, while faded spot bars represent TE family counts for TE families unique to the focal genome assembly. TE classifications are indicated by the key. Numbers at the ends of bars indicate total TE family count within each genome assembly. **ALT TEXT:** Graphs and data on transposable element content annotated among 10 fungi, with subfigures labelled from a to d, illustrating statistical analyses and transposable element variation.

At deep phylogenetic scales, TE content is expected to have almost completely diverged. This is reflected in the *de novo* curation of the fungal dataset producing nearly exclusively TE families unique to a single genome (Figure 4). In total, 2,799 TE families were curated into the shared library spanning the fungal datasets and, contrasting both analyses at smaller scales, the majority of TE families (65.0%, 1,697 / 2,611 families annotated in at least one assembly) are annotated in only a single genome assembly, whilst a very small number of families are shared among all ten genome assemblies (0.50%, 13 / 2,611 families annotated in at least one assembly) (Table S5, Supplementary File 3).

TE coverage is highly variable among fungi, ranging from 5.58% of total genome size in *Paramicrosporidium saccamoebae* (Cryptomycota) to 77.23% in *Golovinomyces orontii* (Ascomycota). The proportion of classified TE content is higher in Ascomycetes and Basidiomycetes, consistent with the bias in genomic analysis towards these phyla. Reflecting well-described patterns in other lineages (Feschotte and Pritham 2007; Canapa et al. 2015; Bourque et al. 2018), we find a significant relationship between TE abundance and genome size (linear regression, F_1,8_=94.25, p < 0.001, adjusted R^2^=0.912), highlighting TEs as a major contributor to genome size variation among these fungal lineages (Figure 4).

At this scale, variation in TE dynamics among lineages become more apparent. For example, among the two Ascomycete lineages, LTR retrotransposons dominate the genome of *Ophiocordyceps sinensis*, contributing 81.8% of total TE content (70.9 Mb), whereas LINEs dominate the genome of *G. orontii*, contributing 48.1% of total TE content (78.4 Mb). Whilst LTR retrotransposons dominate the TE landscapes of Ascomycota and Basidiomycota, DNA transposons are the most abundant classified TEs in *Pinnaticoemansia coranantispora* (Zoopagomycota). Within *P. coranantispora*, DNA TE activity appears to be recent and potentially ongoing, with the majority of DNA TEs showing Kimura 2-parameter distances < 0.1 (Supplementary Figure 1). However, considering the most recent TE activity, represented by a Kimura 2-parameter distance of 0.0, there is a noticeable increase in LINE activity, potentially indicating the start of a new burst of TE activity that could act as a source of genomic novelty, although deeper analyses at the population-level are required to unveil the potential impacts of this apparent recent and ongoing TE activity.

Given the deep divergence among species and the limited shared TE content, substantial differences in the relative distribution of TE copies across divergence classes are expected. In *Allomyces macrogynus*, *Dissophora decumbens*, *G. orontii*, *Piromyces* sp., and *T. hemisulphureum*, TE landscapes show pronounced peaks at very low Kimura 2-parameter divergence (0.01–0.00) from consensus sequences, consistent with recent TE activity or accumulation within these lineages. Conversely, peaks in *O. sinensis* and *P. coronantispora* occur at greater divergence (0.06–0.07), suggesting that major periods of TE activity occurred further in the past and that very recent accumulation is comparatively limited (Supplementary Figure 1). Such patterns could reflect the subsequent establishment of host defences against active TEs, or the loss and degradation of active TE copies through selection and drift. Whereas these species generally show pronounced peaks accompanied by long tails, the landscapes of *N. parisii*, *Paramicrosporidium saccamoebae*, and *Rhizoctonia solani* are characterised by an accumulation of older, more divergent elements. These profiles may reflect historical periods of TE activity followed by reduced activity and retention of degraded copies within the genome. However, these landscapes should be interpreted cautiously as genome assembly quality varies substantially across the fungal dataset. Incomplete or fragmented assemblies may affect TE detection and the representation of TE copies and could therefore contribute to apparent differences in TE landscape shape.

## Discussion

Here, we introduce Earl Grey ParTEA, a framework for scalable and consistent comparative analysis of TEs across multiple genome assemblies. Our applications show that progress beyond single-genome curated TE libraries is critical to capture the complexity of TE diversity. In *A. thaliana*, TE diversity continued to increase with the addition of genome assemblies, indicating that even a panel of ten genomes does not fully capture the TE family diversity present within the species. Substantial amounts of TE content were also restricted to individual assemblies, highlighting marked variation in TE repertoires among closely related genomes. Across the fungal dataset, by contrast, very little TE content was shared among phyla, accompanied by large differences in overall TE abundance and composition. Together, these analyses illustrate how a shared, non-redundant TE library provide a robust framework for investigating TE diversity and distribution across evolutionary scales.

ParTEA produces resampling curves to assess the saturation of TE family discovery. We show that from within species to cross-phylum datasets, saturation is not reached, reinforcing the idea that characterising TE diversity from a single reference genome is misleading. Reaching saturation is not a trivial task and requires both adequate genome sampling strategies but also significant compute resources. Efficient implementations of compute tasks such as the ParTEA snakemake pipeline make such analyses feasible even on moderately sized compute infrastructure.

A considerable proportion of the TE diversity detected in our analyses was assembly specific. These families should, however, be interpreted as unique relative to the sampled genome panel rather than as evidence of *de novo* family origin. Their absence from other assemblies may reflect true absence, insufficient sequence conservation for detection, or limitations in genome assembly and annotation. At the kingdom-wide scale, the near absence of shared TE families across the fungal dataset is consistent with extensive turnover of TE repertoires over deep evolutionary time (Chalopin et al. 2015; Wells and Feschotte 2020; Baril and Hayward 2022), while also illustrating the increasing difficulty of recognising homologous elements as sequence divergence accumulates. Together, these results highlight the highly dynamic and lineage-specific nature of TE repertoires and the importance of using TE resources that are appropriate to the evolutionary scale and taxa under investigation.

The principal methodological advantage of ParTEA is that comparative information is incorporated during TE library construction and annotation rather than being introduced only after independent genome-wise analyses. A single non-redundant library is constructed from TE sequences identified across the dataset and then used to annotate all input genomes, providing a common basis for comparing TE family presence and genome coverage. This approach reduces differences arising from genome-specific TE libraries while enabling direct identification of shared and assembly-specific TE content. Combined with parallelised processing and integrated quality-control summaries, ParTEA therefore provides a standardised workflow for comparative TE analysis as increasingly large genome panels become available.

The integrated BUSCO and phylogenomic analyses provide a useful framework for assessing assembly completeness and organising comparative outputs, but the fungal dataset also highlights limitations of this approach. BUSCO completeness varied substantially among assemblies, from 10.9% to 98.1%, and the resulting phylogeny included placements inconsistent with the expected relationships among major fungal lineages. The misplacement of *N. parisii* illustrates how uneven representation of single-copy orthologues can affect lightweight phylogenetic inference when genome quality varies substantially. The ParTEA phylogeny should therefore primarily be viewed as a tool for organising genome-level summaries rather than as a substitute for dedicated phylogenomic analyses. Where evolutionary relationships are central to interpretation, independently inferred and validated species trees should be preferred.

The interpretation of shared and unique TE content is necessarily dependent on how TE families are delineated. ParTEA uses sequence clustering to provide a consistent operational definition of family membership across the input assemblies, but sequence similarity is only an approximation of evolutionary relatedness and may over-merge closely related elements or separate highly diverged members of the same lineage. This limitation is particularly relevant for datasets spanning large evolutionary distances, where homologous elements may no longer be recognisable by sequence similarity alone. Consequently, shared and unique family counts should be interpreted in the context of the clustering parameters used rather than as absolute measures of TE family diversity, and these parameters should be reported to facilitate comparison among studies.

The substantial contribution of unclassified sequences to the unique TE content identified in the *A. thaliana* dataset further highlights both the value and limitations of *de novo* TE discovery. A mean of 6.04 Mb of unique TE sequence per assembly could not be assigned to currently recognised TE classes, potentially reflecting limitations in existing TE reference resources, rapid sequence divergence, incomplete knowledge of TE diversity, or difficulties in classifying unusual elements. Rather than indicating recent TE origin, these sequences provide a resource for identifying previously under-characterised components of the mobilome. More broadly, the increasing availability of genome panels means that comparative *de novo* approaches such as ParTEA may contribute not only to consistent annotation, but also to the discovery and characterisation of TE diversity that is absent from current reference resources.

Despite the rapid expansion of genomic resources, comprehensive and consistently curated TE reference libraries remain unavailable for many taxa. Manual curation remains important for producing high-quality TE models, particularly when novel or highly divergent elements are encountered, but it is labour-intensive and difficult to scale across large numbers of genomes. ParTEA can help bridge this gap by producing extended consensus sequences across a genome panel, providing a starting point for subsequent manual curation and reducing the need to rediscover candidate elements independently for each genome (Confais et al. 2026). This approach may also facilitate the development of community resources in which computationally generated libraries are progressively refined through targeted curation. The recently developed MycoMobilome (Baril and Croll 2026) resource illustrates this possibility, using thousands of fungal genomes to generate a large non-redundant collection of TE consensus sequences that can subsequently serve as a foundation for lineage-specific annotation and curation.

## Future Developments

ParTEA provides a foundation for scalable and reproducible comparative TE annotation, but several aspects of the framework could be further developed. Future iterations could improve the completeness and curation of TE libraries, enhance detection of rare or structurally complex elements, and incorporate additional approaches for TE classification and quality control. These developments would complement the current repeat-first workflow while extending its utility across increasingly large and diverse genome datasets.

A remaining limitation of the current ParTEA workflow is the detection of rare or low-copy TE families. RepeatModeler2 and related repeat-first approaches exploit the repetitive nature of TEs, and elements represented by few genomic copies may consequently be more difficult to identify. The multi-genome design of ParTEA can partially alleviate this limitation when a rare family occurs at higher copy number in another assembly within the panel, as observed in our *Drosophila* panel in this study and during the construction of a comprehensive TE library for *Zymoseptoria tritici* (Baril and Croll 2023). However, this remains dependent on the family being sufficiently represented in at least one input genome. Pangenome-based approaches provide a complementary strategy by identifying TE-associated insertion polymorphisms directly rather than relying on repeat abundance. For example, Pantera (Sierra and Durbin 2024) and GraffiTE (Groza et al. 2024) use polymorphic sequences from pangenome graphs to identify TE families and have been shown to recover large, low-copy elements that can be missed by conventional de novo approaches. Incorporating such approaches into ParTEA could improve sensitivity to rare and structurally complex TEs, although their dependence on closely related genomes and greater computational requirements would limit their applicability to the deepest comparative scales supported by ParTEA.

The TE libraries generated by ParTEA could also be subjected to additional post-discovery curation. De novo repeat discovery can occasionally incorporate multicopy host genes or other non-TE repetitive sequences into TE libraries, while genuine TEs may remain difficult to classify when they are highly diverged or lack sufficient similarity to existing reference models. Future versions of ParTEA could therefore incorporate screening against host gene annotations and other genomic resources to flag likely gene-derived sequences. In parallel, analyses of conserved TE-associated domains could improve classification where sequence-homology approaches provide limited information. Recent tools such as MCHelper (Orozco-Arias et al. 2024) demonstrate the feasibility of automating several aspects of TE-library curation and classification across diverse eukaryotic genomes. Such approaches could allow ParTEA to move beyond automated consensus generation towards more systematic computational curation of TE libraries.

## Conclusions

Overall, Earl Grey ParTEA extends automated TE annotation from individual genomes to comparative genome panels, providing a unified framework for TE discovery, annotation, and cross-genome comparison. The analyses presented here demonstrate that broader genome sampling can reveal substantial additional TE diversity within species while also exposing extensive differences in TE repertoire and composition across evolutionary scales. By integrating dataset-wide TE discovery with consistent annotation and comparative summaries, ParTEA provides a practical framework for investigating TE diversity as genomic resources continue to expand.

## Availability and Requirements

Project name: Earl Grey ParTEA

Project home page: https://github.com/TobyBaril/EarlGreyParTEA

Operating systems: Linux-based systems (e.g. Ubuntu)

Programming Language: Pipeline implemented in Snakemake, including software coded in Python, R, Bash, and Perl.

Other requirements: A conda package manager (e.g. Anaconda3, Miniforge, Micromamba, Pixi)

License: Open Software License v 2.1

Restrictions for non-academic users: none. Some dependencies may require licenses for use by non-academic users.

## Acknowledgements

We thank Dr. Olivier Coen for discussions on implementation and unit testing of Earl Grey ParTEA. We thank users of early iterations of the software for constructive feedback and suggestions.

## Author Contributions

T.B. conceived the study and developed Earl Grey ParTEA, performed analyses, produced figures, and drafted the manuscript. A.F. developed the initial Snakemake framework and assisted in the integration of Earl Grey workflows into Earl Grey ParTEA. D.C. conceived and coordinated the study, participated in writing the manuscript, and provided funding. All authors read and approved the final manuscript.

## Data Availability

All data generated or analysed during this study are included in this published article and its supplementary information files, which are hosted on Zenodo under the DOI: (10.5281/zenodo.23955171).

